# Data-driven modeling of group formation in the fission-fusion dynamics of Bechstein’s bats

**DOI:** 10.1101/862219

**Authors:** Nicolas Perony, Gerald Kerth, Frank Schweitzer

## Abstract

Communal roosting in Bechstein’s bat colonies is characterized by the formation of several groups that use different day roosts and that regularly dissolve and re-merge (fission-fusion dynamics). Analyzing data from two colonies of different size over many years, we find that (i) the number of days bats stay in the same roost before changing follows an exponential distribution that is independent of the colony size, and (ii) the number and size of groups bats formed for roosting depend on the size of the colony such that above a critical colony size two to six groups of different sizes are formed. To model these two observations, we propose an agent-based model in which agents make their decisions about roosts based on both random and social influences. For the latter, they copy the roost preference of another agent which models the transfer of the respective information. Our model is able to reproduce both the distribution of stay length in the same roost and the emergence of groups of different sizes dependent on the colony size. Moreover, we are able to predict the critical system size at which the formation of different groups emerges without global coordination. We further comment on dynamics that bridge the roosting decisions on short time scale (less than one day) with the social structures observed at long time scales (more than one year).

## 1 Introduction

The idea that “more is different” (Anderson, 1972) has become a common paradigm to describe a system whose behaviour changes qualitatively when the number of its elements increases. As emphasised by Cavagna and Giardina (2010), it is also an interesting perspective to look at animal groups. These groups vary widely in size and stability, from small social groups with stable individual composition as in cooperatively breeding mammals to vast aggregations such as kilometre-long fish shoals containing tens of millions of individuals (e.g Krause and Ruxton (2002)). Large variations in group size can also be observed within the same species. Increasing group sizes have been shown to foster division of labour (Gautrais *et al*., 2002; Jeanson *et al*., 2007), transitions from disorder to order (Buhl *et al*., 2006), or the accuracy of group decisions (Couzin, 2009). Generally, these questions are linked to the topic of optimal group size in animal populations (Krause and Ruxton, 2002).

Here we concentrate on the Bechstein’s bat (*Myotis bechsteinii*), a species where the females form maternity colonies with high fission-fusion dynamics in summer, during the breeding season (for a detailed species description, see Kerth, 2006). *Fission-fusion dynamics* refers to the regular splitting into and merging of *groups* within larger social entities such as colonies in case of bats (Aureli *et al*., 2008; Couzin and Laidre, 2009; Sueur *et al*., 2011). Previous studies (Couzin and Krause, 2003) showed that fission-fusion dynamics may result from simple association mechanisms, and often produce right-skewed group size distributions, with many small groups and very few large ones.

Compared to the number of empirical studies on fission-fusion dynamics (see e.g. Couzin and Laidre, 2009; Aureli *et al*., 2008; Van Horn *et al*., 2007; Ramos-Fernández *et al*., 2006; Willis and Brigham, 2004) modeling approaches are less developed. They can be divided into statistical models and generative models. Statistical models, for instance regression models, aim to *infer* from available data the influences that govern the observed dynamics. For example, the frequency of fission and fusion events in reindeer was predicted based on the observed variation in group sizes (Body *et al*., 2015). An advanced statistical model, the hierarchical Bayesian model, was used to disentangle the influence of other individuals (action, sex) on the individual fission and fusion decision of spider monkeys (Ramos-Fernández and Morales, 2014).

The evaluation of statistical models is usually restricted to comparing the statistical performance of model variants with and without certain influences. This allows to estimate the impact of these influences on explaining the data, but it gives no insights into the interaction dynamics or decision rules of individuals. This methodological limitation is tacked by generative models, for instance agent-based models. These propose rules, e.g for interactions or decisions, and then test to what extent such assumptions are compatible with an observed behavior, either on the individual or the systemic level. An agent-based model of the seasonal fission-fusion dynamics in red-capped mangabeys, for example, was able to reproduce observed patterns of travel distance (Dolado *et al*., 2017). Also, the impact of individual compromises between nutritional needs and social interactions on the social network between individuals and a possible irreversible fission was simulated (Sueur and Maire, 2014) with an agent-base model. These type of models also allow to test the impact of certain parameters, for example split-rates (Conradt and Roper, 2000; Nair *et al*., 2019), on the principal outcome of the fission-fusion dynamics. In most cases, however, the assumed rules cannot be directly matched to available observations. Therefore, agent-based models provide a way to develop hypotheses about unobserved behavior which can later be addressed in subsequent research. We follow this approach in our paper, with a specific focus on bats.

In bats, most species are social and form groups of variable size and composition, and many of the colonies display fission-fusion behaviour (Kerth, 2008). Female Bechstein’s bats profit from group formation while roosting as they obtain thermoregulatory benefits from clustering (Pretzlaff *et al*., 2010; Küpper *et al*., 2016). Roosting groups are the result of collective decisionmaking about communal roosts (Kerth *et al*., 2006; Fleischmann *et al*., 2013). As Bechstein’s bats forage separately or in pairs at night and do not come back to the roosting area all at the same time (Melber *et al*., 2013), information available to each individual about the roosting preferences of the other colony members is limited. Field experiments have shown that colony members exchange some information about the location of suitable roosts (Kerth and Reckardt, 2003). In such an environment, the existing literature (Conradt and Roper, 2003, 2005, 2007) suggests that the decision mechanisms involved in the choice of a roosting site may be self-organized.

In this paper, we describe such a self-organized decision-making process in Bechstein’s bats. To gain empirical insights, we analyze data on the daily roosting behaviour of individuals from two colonies (see Section 2.1 and Kerth *et al*. (2011) for a full description of the data). These data report about the outcome of the collective decision process, in terms of groups formed for communal roosting. But the rules bats follow to make these decisions about their roosts are unknown. Therefore, in this paper we propose such rules which take individual preferences and the information transfer between conspecifics into account.

In addition, we focus on the *group sizes* that result from such collective decisions. The question of how groups form and how their size is regulated is still a very topical one, as accurate studies of the temporal dynamics of an animal group are rarely paired with theoretical justifications derived from robust models (Sumpter, 2010). The study and characterisation of group sizes is an even larger challenge when the animals undergo a fission-fusion dynamics as in the case of Bechstein’s bats.

Our main interest is to explain how the formation of roosting groups is influenced by the size of the colonies. If “more is different”, we should expect the emergence of specific collective behavior once colonies have reached a critical size. As we report in the empirical findings, the formation of roosting groups of different sizes dependent on the size of the colony is such an emergent phenomenon. We demonstrate that this transition in the behavior of the colony can be well explained by our model. But we also show that our model is able to reproduce the distribution of durations bats spend in the same roost before switching to another roost.

Eventually, in this paper we also investigate how the formation of groups as part of the fission-fusion dynamics can be related to the emergence of long-term social structures, such as the existence of communities in social networks of large colonies of Bechstein’s bats (Kerth *et al*., 2011).

## 2 Methods and Data

### 2.1 Available data and subsequent measures

For our model validation, we have data available for two different colonies of Bechstein bats (*Myotis bechsteinii*), a larger one denoted by GB2 with 34-46 individuals, and a smaller one denoted by BS with 11-18 individuals (Kerth *et al*., 2011; Baigger *et al*., 2013). Both colonies were observed over many years, from 2004 to 2010 for GB2 and from 2004 to 2008 for BS. All bats in both colonies have been individually marked with PIT-tags in their first year of life (Kerth, 2006), i.e. they can be identified by these tags over years.

Although Bechstein’s bats forage separately or in pairs during the night, they have to roost together during the day, to benefit from social thermoregulation (Pretzlaff *et al*., 2010; Küpper *et al*., 2016). I.e., they form small groups that occupy a “bat box” for some days, but then have to change their box because of the need to avoid parasites that accumulate in the boxes (Reckardt and Kerth, 2006) and to find optimal roosting temperatures that are weather dependent (Kerth *et al*., 2001b). One to six of such roosting groups per colony are formed, and their composition can alter every day. These groups can choose from about 150 bat boxes that were placed in the home range of the two colonies (Kerth *et al*., 2011). Only about 50 different boxes (out of 150 available in both colonies, together) are being occupied by the groups in each season, and members of different colonies do not roost together. Each of these boxes is equipped with an antenna that is connected to an automatic PIT-tag reader that stores PIT-tag numbers, times, and dates of each bat entering the box. This way, from 2004 onward, for the breeding season between April and September, we have daily data about the presence of individual bats in the respective box.

To formalize the information available from the data, we first introduce three different *time scales*. The longest time scale, *y*, is measured in *years*, or seasons. One season consists of about 200 days, during which information about the colony becomes available. This is the time scale at which long-lasting social structures of the colony become visible, such as communities Kerth *et al*. (2011). We will come back on this in Sect. 3.3.

The intermediate time scale, *t*, is measured in *days*, i.e. it is also a discrete scale. On this time scale, the so-called fission-fusion dynamics becomes important. Bats form groups for communal roosting (fusion), however these groups are not stable over a long time and dissolve mostly during one to two days (fission). On time scale *t* bats decide about their (daily) roost, for which we have information available. The automatic reading lead to 6655 individual roosting records for BS and 13845 for GB2. About 97% of the bats passing the antenna in the box entrance could be identified (Kerth and Reckardt, 2003).

This registration allows us to subsequently calculate *pairwise roosting associations* for each colony. If *r* ∈ {1,…, *m*} is the discrete number of the box, then *r_i_*(*t*) tells us that bat *i* has roosted in box *r* at day *t*. The Kronecker delta *δ_ij_*(*t*) then indicates whether two individual bats *i* and *j* have roosted together at that particular day *t. δ_ij_*(*t*) = 1 if *r_i_*(*t*) = *r_j_*(*t*), and *δ_ij_*(*t*) = 0 otherwise. Aggregating over time for a fixed pair of individuals *i, j* tells us how often these two bats have roosted together. If the latter is normalized to the number of days both of these individuals have been observed in the area, it yields the *I_ij_* index (Kerth and König, 1999).

The shortest time scale, *τ*, is much shorter than one day and could be measured e.g. in *minutes*. In comparison with the time scale *t*, we can treat this time scale *τ* as (quasi) *continuous*. This is the time scale at which bats *exchange information* about suitable roost sites and *decide* where to roost. We see this communal roosting as the outcome of a *collective decisions process*, for which we can observe the *result* (on the time scale *t*), but do not know the *rules* which generate the observed outcome on the time scale *τ*. To infer a set of possible decision rules that are compatible with this outcome is precisely the aim of our paper. This requires us to model the so-called *swarming phase* more explicitly, during which bats exchange the above information. The swarming phase describes the aggregation of bats that fly around a box at dawn before they will eventually use it as day roost (Nad’o and Kaňuch, 2015). During this swarming phase the bats presumable make their collective decisions about where to communally roost during the day. Before we come to this, we need to take a closer look at characteristic features of the roosting data, obtained at time scale *t*.

### 2.2 Distribution of duration of stay in a box

As we have already mentioned, Bechstein bats have an incentive to switch nest boxes. Hence, the first question is about their average roosting duration in a given box. To calculate this duration *T*, measured in days, we have to compare, for each bat, their daily roosting locations at consecutive days. This results in a time series of values of *T_i_* for each individual bat. To characterize the colony, we have to determine the distribution *P*(*T*) from the histogram of all *T* values for all bats from the same colony. The result is shown in Fig. 1. We find, for both colonies, the same exponential distribution *P*(*T*) ∝ *e^−αT^* with almost the same values of *α*. The statistical details are given in the Appendix.

**Figure 1:**
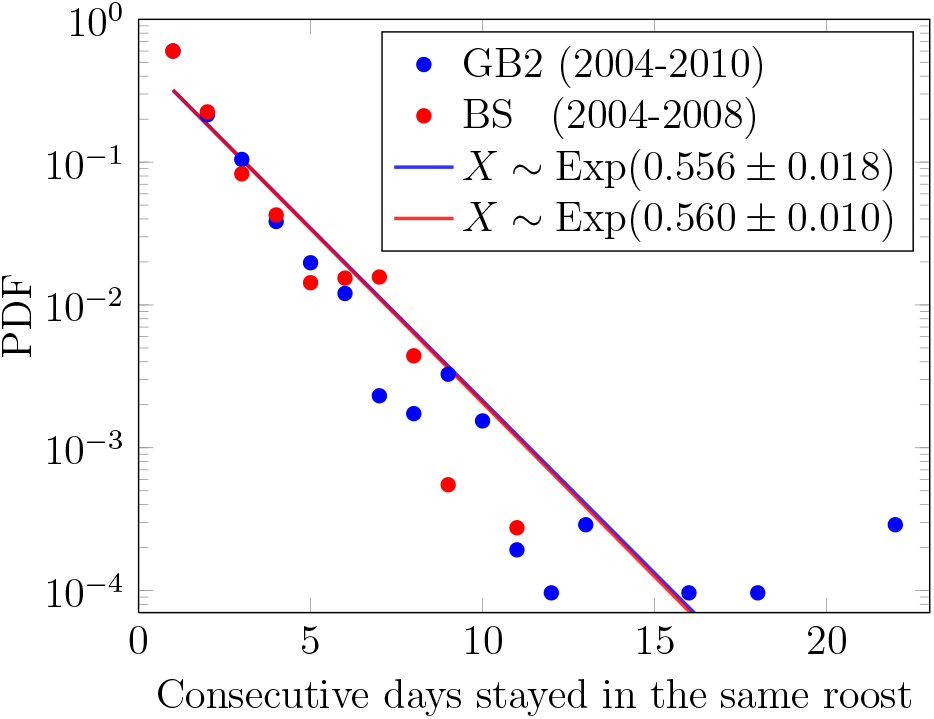
Probability density function (PDF) of the duration of stay for a bat in a given box. The solid lines are fits for both colonies to exponential distributions, with the scale and its 95% confidence interval indicated in the legend.

It is remarkable that the distribution of duration to stay in the same box does not differ for the two colonies of very different sizes, GB2 being about twice as large as BS. This indicates that biological reasons that are largely independent of total colony size such as parasite infestation in the roost (Reckardt and Kerth, 2006) or the roosts’ micro-climate in relation to weather conditions (Kerth *et al*., 2001b) determine the duration to use the box. In both colonies, we observed that bats changed boxes on average about every two days, because the mean period is the inverse of the distribution’s rate α. This is in line with values found in previous studies on Bechstein’s bats (Kerth and König, 1999; Kerth *et al*., 2011; Fleischmann and Kerth, 2014) and in maternity colonies of other species of forest-dwelling bats (e.g. O’Donnell and Sedgeley (1999); Willis and Brigham (2004)).

### 2.3 Distribution of group sizes

In a second step, we focus on the size of the groups that roost together in one box. Both the number and the maximal size of these groups depend on the size of the colony, which is very different for GB2 and BS. Hence, we have to distinguish between three different levels:

- *N*(*y*) is the size of the *colony*, which can vary from year to year, as Figure 2 shows, but is assumed as fixed for a given year *y* because of the high individual stability of the colonies and the very low mortality of the bats during summer (Fleischer *et al*., 2017).
- Because of the fission-fusion dynamics, each colony is composed of *groups* of different sizes, *n_k_*(*t*), where *k* is a group index and *n_k_*(*t*) is the size of the group *k* at a particular day *t*. The boundary condition 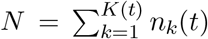 has to be fulfilled for each day. The total number of groups, *K*(*t*), is not a constant, but can vary on a daily scale. For comparison of colonies with different sizes, we introduce the *relative group size, *x*_k_*(*t*) = *n_k_*(*t*)/*N*, with ∑_*k*_ *x_k_*(*t*) = 1
- On the third level, we have *individuals i* with *i* = 1,…,*N* that compose the different groups. The composition of the groups can also vary day by day. Sometimes a group can only consist of a single individual that roosts alone, at other times all individuals may form a single group of the size of the colony. I.e., *x_k_*(*t*) can vary between 1/*N* and 1 in the extreme cases, which also impacts the total number of groups per day, *K*(*t*).

**Figure 2:**
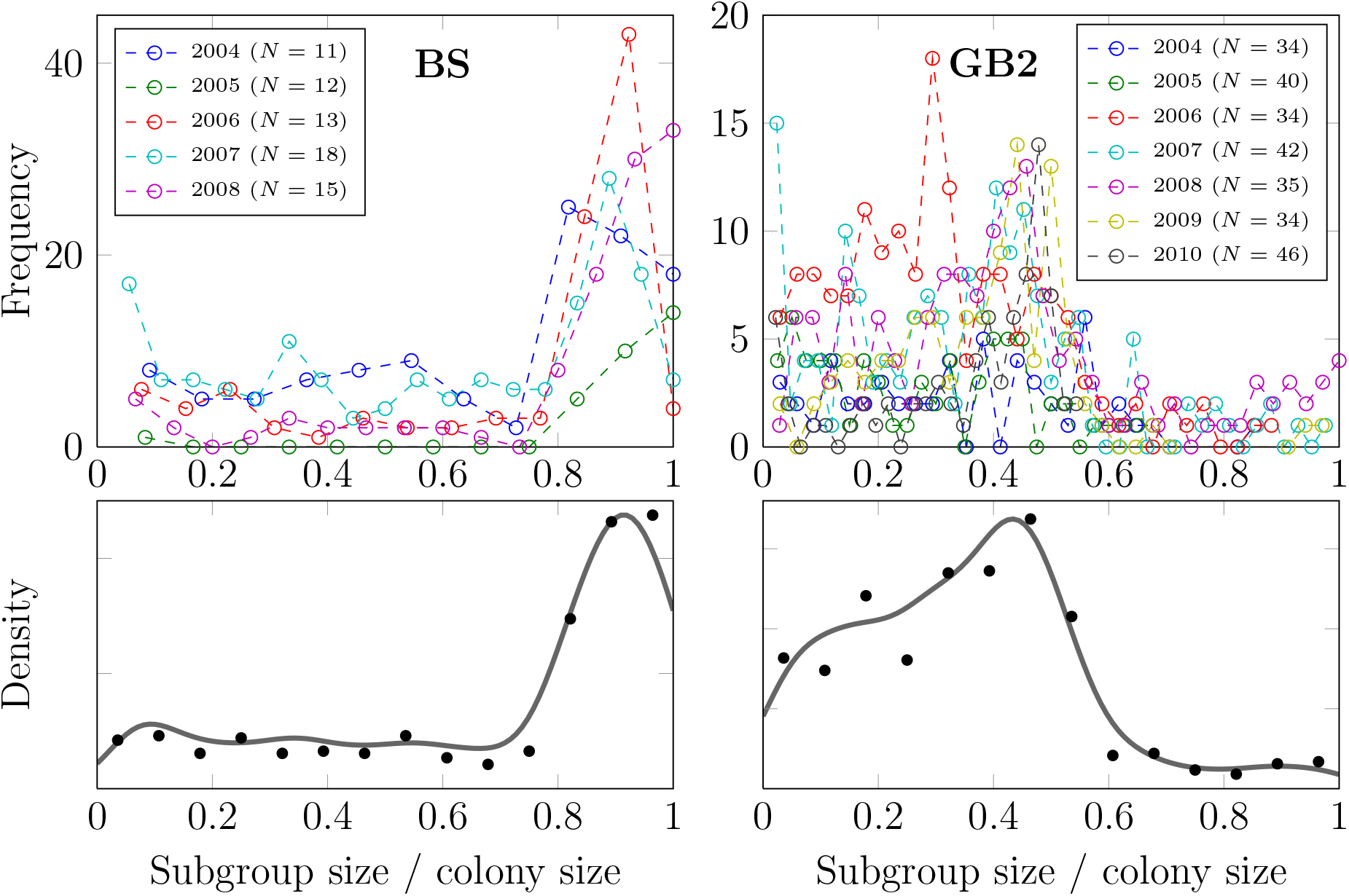
Distribution of relative group sizes *x_k_* in colony BS (left panels) and GB2 (right panels). Measures are presented for both colonies during all years of the study (top panels) and aggregated over all years (bottom panels). The legend in the top panels indicates the size of each colony on a given year. The aggregated density values (bottom panels, black dots) were smoothed using a Gaussian kernel density estimate (solid thick lines) to give a visual indication of the shape of the distribution.

In Figure 2, we calculate, for each colony separately, how often relative group sizes *x_k_*(*t*) have been observed in a given year. To highlight the differences, we have then calculated from these frequencies, aggregated over all years, the distribution *P*(*x*) for the two colonies.

Comparing the two distributions *P*(*x*), we already note that they vary remarkably. The small colony BS displayed a very cohesive behaviour, i.e. all individuals mostly roosted together. The large colony GB2, on the other hand, showed the formation of groups of a size smaller or approximately equal to one half of the colony size. Only in very few occasions the whole colony was roosting together. There are two remarkable observations from Figure 2:

i. Bechstein’s bats indeed make collective decisions when it comes to roost choice, otherwise the formation of groups and the coordinated behavior would not be observed. This confirms previous findings (Kerth *et al*., 2006; Fleischmann *et al*., 2013).
ii. Comparing the small and the large colony, we argue that there is a critical colony size *N_crit_* above which the formation of groups becomes very likely. The graphs indicate that this critical size is roughly around 18, because the smaller colony most often roosted together, whereas the larger colony most often formed groups of size 1/3 to 1/2 of colony size.

### 2.4 The need to model social influence

In the following, we develop an agent-based model that aims at reproducing the two mentioned empirical observations, (i) the distribution of consecutive days of staying in the same box, Figure 1, (ii) the size of the groups that roost together, dependent on the colony size, Figure 2.

To start with the exponential distribution of durations *T*, we know that such a distribution can be obtained from assuming a simple Poisson process for bat’s interactions. Specifically, we could consider that agents every day change from their previously occupied roost to another roost with a fixed probability *α*. So, the chance that they stay at the same roost is decaying as 1 − *α*, and then they decide for another box.

One could argue that this decision to leave depends on the available roost sites in the vicinity. Because flying long distances is energetically costly for broad winged bat species (see e.g. Entwistle *et al*., 1996), Bechstein’s bats may prefer to fly rather short distances when switching to another box, thereby minimising their energy expenditure. However, in Bechstein’s bats the foraging area of a bat is typically much larger than its roosting area and thus roost-switching distances may not be relevant to its box choice process (Kerth *et al*., 2001a). Testing for this effect, we found no effect of the previously occupied box on the next occupied box in terms of flying distance between boxes. The statistical details are again given in the Appendix.

This insight then lends evidence to the assumption that bats choose *randomly* among the available roost boxes. Hence, one could assume that the whole process of leaving a box and choosing another box can be modeled as a random process, where the fixed probability α decides *when* to leave.

With such an assumption, we correctly reproduce the distribution *P*(*T*) observed in Figure 1, independent of the system size. But, as a consequence from this assumption, we *also* get an exponential distribution of *group sizes*, because agents find themselves together only at *random*. There is evidence for such an exponential distribution of group sizes for *other systems* (Okubo, 1986; Bonabeau and Dagorn, 1995; Flierl *et al*., 1999; Bonabeau *et al*., 1999; Couzin and Krause, 2003), but *not* for *our* system of Bechstein’s bats, where the distribution of group sizes is very different from an exponential distribution. As Figure 2 shows, in our case the distribution is *not* right-skewed as it is the case for an exponential distribution. Further, our distribution also *changes* based on the system size, i.e., the number of bats.

Hence, from these considerations we can conclude that it needs another mechanism in the model to correctly account for the way bats communicate about their roosting intention, and form these groups. Therefore, in addition to the *random influence* already assumed for reproducing the exponential distribution of durations, *P*(*T*), we have to add *social influence*. Only this will allow agents to copy the roosting intention (or preference) of other agents, as needed to reflect the collective decision.

### 2.5 Modeling the swarming phase

As already argued, social influence is exerted at a time scale *τ* shorter than the scale *t*, specifically during the *swarming phase* (at dawn), in which Bechstein’s bats aggregate in flying around potential roost with the opportunity of signalling their preferences for certain day roosts to other colony members. We assume that, when the swarming phase starts, each agent has a roosting preference *r_i_*(*τ*), where *r* ∈ {1,…, *m*} is the identity (number) of the preferred box *r*. The start value for *r_i_*(*τ*) is the roost number from last day. This preference is not fixed, but can change during the swarming phase. In our model we assume that this dynamics is governed by two different processes: (i) a *random change*, which is modeled again by a Poisson process with a rate λ (equal for all agents), and (ii) the *social influence*, which causes an agent *i* to change its roost preference *r_i_* to the *r_j_* of another agent *j* at a rate *γ*.

So, basically these two processes compete: agent *i* picks a new preferred roost either *randomly* or takes the roost preference of other bats into account (collective decision-making). The latter means that agent *i amplifies* the preference *r_j_* for a given box by *copying* the respective decision from agent *j*. Which of these processes dominate, depends on the ratio λ/*γ*, which will be determined later during the model *calibration*.

In order to decide when a roosting preference *r_i_*(*τ*) is finalized, i.e. does not change further, we could set an arbitrary time after which the swarming phase is finished. But this would introduce a strict cutoff in the model that can hardly be justified. It further increases the influence of noise on the dynamics, because it is rather arbitrary which preferences agents have at a fixed point in time. To avoid such artifacts, we model a progressive decision process in which agents switch one by one to the decided state at a rate *ξ*. We can set *ξ* = 1 for simplicity because both λ and *γ* are defined relative to *ξ*. Implementing the decision process this way allows a rich dynamics in which agents finalize their preferences at different times. This allows agents who have already decided about their roost to still influence agents who have not yet decided where to roost.

## 3 Results

### 3.1 Model calibration

To obtain results, we need to calibrate the two parameters introduced, (i) the rate λ at which agents *randomly change* their preferences for a roost site during the swarming phase, and (ii) the rate *γ* at which agents copy the preferences of other agents. For this calibration, we use the empirical finding *P*(*T*) of Figure 1 that demonstrates the outcome of the *combined processes* which jointly determine when agents change their current roost.

Our model generates for each agent a sequence of roost sites *r_i_*(*t*) used at consecutive days. From this time series, we can determine the sequence of durations 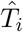 that agent *i* stays in a given box before changing to another box. We deem our model correct, if it is able to reproduce the empirical finding *P*(*T*), i.e. if the model-generated distribution 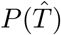 of durations matches the observations. This implies three requirements: (i) 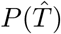 has to be an exponential distribution, which is assured because we have modeled the random change of preferences as a Poisson process. (ii) The characteristic parameter 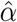 obtained from 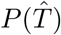 has to match the empirical value *α* = 0.56. (iii) As an additional constraint, we need to make sure that the model-generated distribution 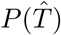 ft also independent of the system size *N*.

These three requirements can be achieved by adjusting the model parameters λ and *γ* such that the match between model and empirics is as good as possible. Specifically, the following two errors have to be *minimized*:

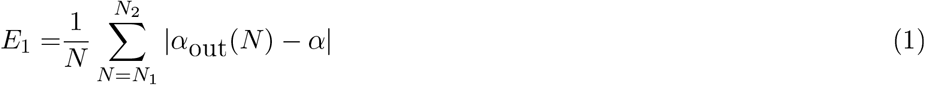

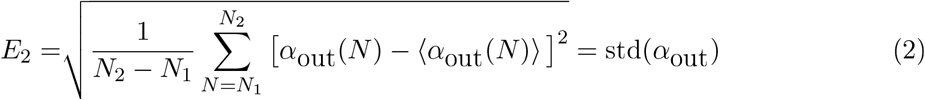

E_1_ measures the difference between *α*_out_(*N*), the model-generated decay value of the exponential distribution, and the empirical value α, which should be as small as possible. The model output *α*_out_(*N*) depends on the colony size *N*, which we have varied in discrete steps between {*N*_1_… *N*_2_}. Practically, we have chosen *N*_1_ = 10 and *N*_2_ = 50 because this is the typical minimum and maximum colony size, respectively (Kerth, 2008). For each value of *N*, we ran 10’000 simulations of the model, hence *α*_out_(*N*) already gives the average over 10’000 simulations.

E_2_ is the standard deviation of the distribution of all *α*_out_(*N*) obtained in the range *N* ∈ {*N*_1_ … *N*_2_}. This error should be minimized during the calibration because we want the model output be independent of the system size *N*. 〈*α*_out_(*N*)〉 is the mean value of *α*_out_(*N*).

To obtain the pair {λ, *γ*} that best fits the experimental data, we minimised the product of the squared error of the exponential fit by the size-related error, 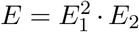. Plots of *E*_1_, *E*_2_ and *E* are shown in Figure 3, We found a minimum of *E* for λ = 0.95 and *γ* = 22.

**Figure 3:**
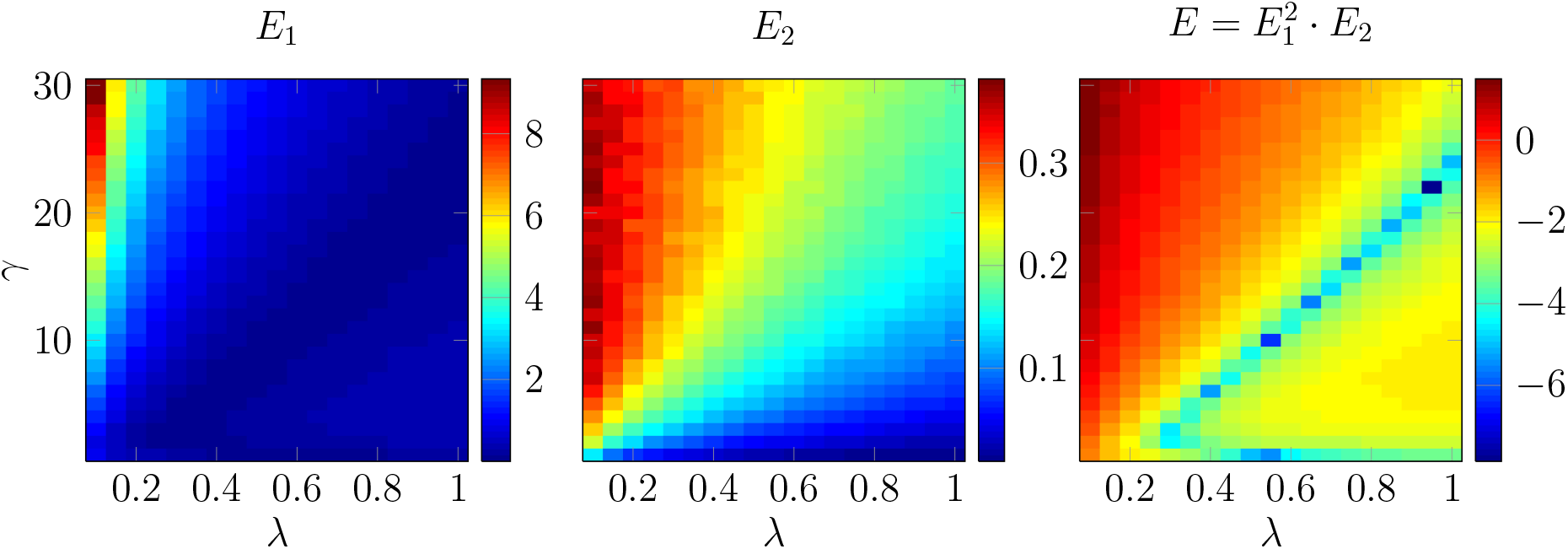
Parameter fitting by minimisation of the model’s total error *E. E*_1_ represents the error of the fit with regard to the experimental data and *E*_2_ the variation of the mean consecutive roosting period with regard to the system size *n* (number of agents), with *n* ∈ {10… 50}. For all plots, the error should be minimised thus less (dark blue) is better; scales are logarithmic.

### 3.2 Distribution of group sizes

The agent-based model outlined above shall now allow us to reproduce, and to understand, the second empirical finding, namely the distribution of the sizes of groups that roost together, dependent on the size of the colony, as shown in Figure 2. Specifically, the small study colony roosted mostly as a single group, while the larger colony roosted in several groups, with the largest roosting group comprising mostly about half of the colony.

However, we do not have observations about the *transition* from one to several groups. Therefore, in a first step we merge the information about the group sizes of the two colonies. Figure 4(left) shows the complete empirical data. The *x*-axis displays the size of the colony, *N*, varying as before between *N*_1_ = 10 and *N*_2_ = 50. The *y*-axis displays compressed information about the size of the groups. The diagonal *y* = *x* shows the *maximum size* of a group for a given size of the colony. If all indiviudals belong to only *one* group, then we should find the observed group size very close to this diagonal. This is indeed the case, as Figure 4(left) shows, but only as long as *N* is below 20. Specifically, we plot the probability of an individual to be part of a group of a given size in terms of a color code. The darker the color, the larger this probability.

**Figure 4:**
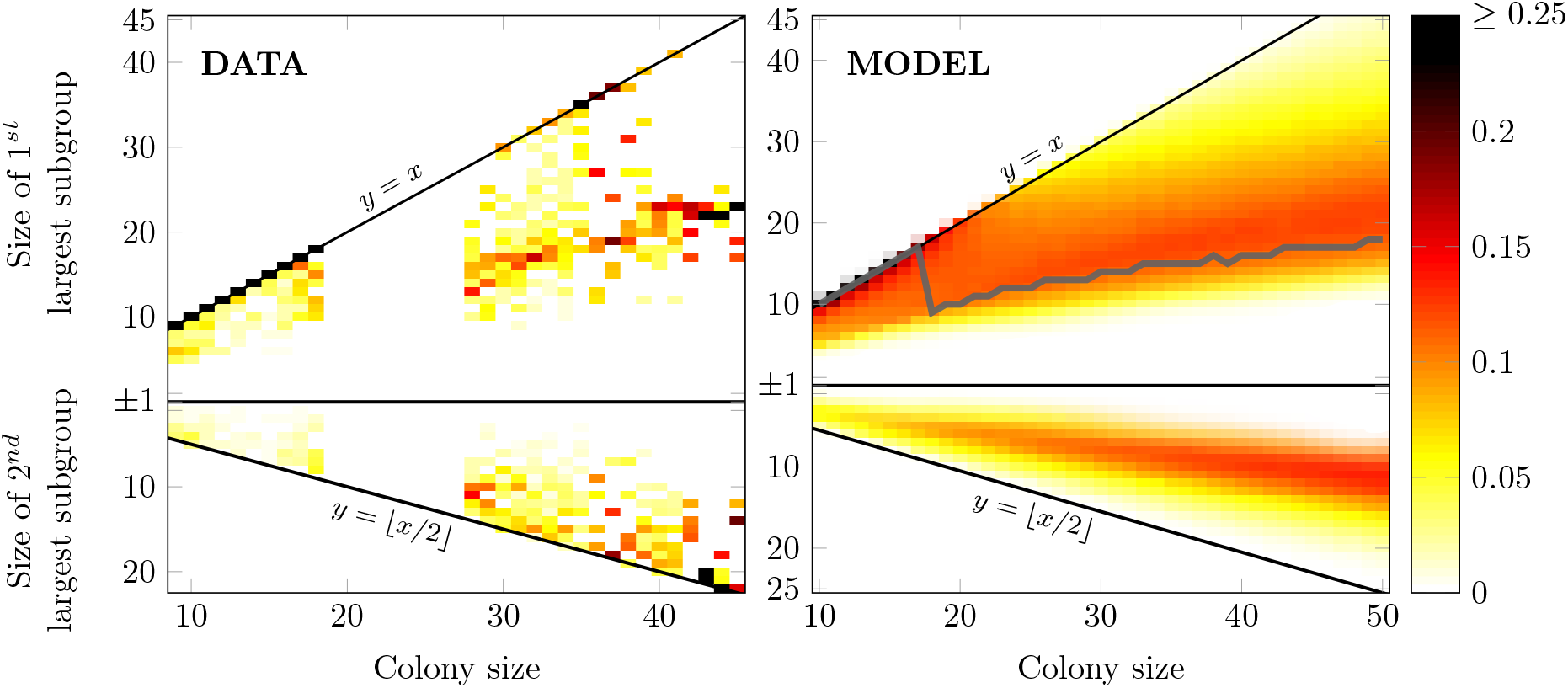
Empirical group sizes (left) and modeled group sizes (right) as a function of colony size. Colors represent the probability for an individual to belong to a group of a certain size. The top part of both figures shows the size of the largest group, the bottom part the size of the second largest group. The line *y* = *x* indicates that there is only one group of the size of the colony. If a second-largest group exists, its size can be maximally *y* = *x*/2 indicated in the lower part. The gray line on the right panel represents the most common largest group size in our simulation results.

For colony sizes *N* between 18 and 26, we do not have any empirical data. But for *N* > 26, we see that the group size quite often differs from the colony size, i.e. most individuals are found in groups of sizes much below the diagonal *y* = *x*. This clearly indicates the formation of groups inside the colony. To better understand how the colony splits into groups of *different sizes*, we have plotted in Figure 4(left) the size of the largest group, *n*_1_(*t*), in the upper part and the size of the second-largest group, *n*_2_(*t*), in the lower part (we have used the group index *k* to rank groups according to their size *n_k_* such th at *k* = 1 refers to the largest, *k* = 2 to the second largest group).

Obviously, the maximum size of *n*_2_(*t*) is bound to *y* = *x*/2, otherwise it would be the largest group. We see that the empirical sizes of the second-largest group are close to this line, but not too close. That means that colonies do not split precisely into two groups of size *N*/2, which would also not be realistic. Instead, we note the formation of the second-largest group with sizes *n*_2_(*t*) comparable to about *N*/3. The sizes of the largest group, *n*_1_(*t*), shown in the upper part, are comparable to *N*/2. This means that, if other groups exist, they can be only of rather small sizes, summing up to about *N*/6.

The right part of Figure 4 shows the same diagram with the results of our calibrated agent-based model. We note that simulations were also performed for colony sizes where no empirical data was available (18 to 26 individuals). The gray line on the right panel shows the most common size obtained for *n*_1_(*t*) in our simulations. This curve displays a sharp drop for a colony size of *N* = 18 individuals. Hence, it marks the transition from a regime, where only *one group* of the size of the colony is observed, to a regime with *multiple groups* of different sizes. The *critical size* of the *colony* for this transition is *N_crit_* = 18.

We note the very good agreement of our model results with the empirical observations. Despite the fact that we only used two parameters λ, *γ* to describe the roosting decision of agents, the model is able to reproduce the findings about group sizes dependent on colony size. This leads us to the conclusion that the underlying dynamics for the agents capture the systemic behavior to a remarkable degree. In particular, the simulations allow us to project the dynamics of the colonies to the unobserved cases. This way, we identified the *critical system size N_crit_* = 18 at which the bifurcation in the systemic behavior, i.e. the transition from a single to a multi group regime, occurs. We further elaborate on this fact in the Discussion.

### 3.3 Including the roosting history

So far, the two rates λ and *γ* have been the same for all agents. This is justified for λ because random influences are considered. For *γ*, however, one would expect that the *social influence* between any two agents also depends on their previous experience together. Hence, instead of a overall rate of social influence, we introduce a pair-specific social influence *γ_ij_*(*t*) that depends on the joint history of agents *i* and *j*. If these agents have roosted together at a particular day *t*, this should increase their mutual social influence by an amount Δ_*γ*_. If, on the other hand, these agents never roost together again, this mutual social influence *γ_ij_*(*t*) should decay over time at a rate *ε*. This can be expressed by the following discrete dynamics:

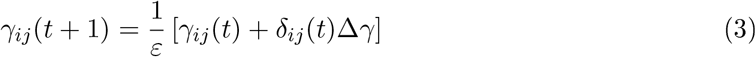

We remind that *δ_ij_*(*t*) is the Kronecker delta, which equals one whenever *r_i_*(*t*) = *r_j_*(*t*), i.e. when agents *i* and *j* roost together at day *t*, and zero otherwise.

The dynamics of Eq. (3) follows the idea of *reinforcement learning*, because a previous joint experience in roosting increases the mutual social influence, which in turn increases the future chances that either agents *i* or *j* copy the roost preference of the other agent. Hence, it describes the formation of social bonds between agents that could also impact the long-term social structures, as discussed below. Mutual social influence that is not maintained, however, will decay over time.

*γ_ij_*(0) denotes the start value of mutual social influence. To set this value, we account for the fact that social influence between individuals of the same group is larger than between individuals of different groups. Hence, initially we create two groups of equal size *N*/2. Within these two groups, we set *γ_ij_* (0) = 0.55 for all individuals in the same group, and between these two groups we set *γ_ij_*(0) = 0.45. Further, we choose Δ_*γ*_ = 0.05 and 1/*ε* = 0.95. The latter describes an exponential decay of the mutual influence, *γ_ij_* (*t*) = *γ_ij_* (0) exp{[(1/*ε*) − 1] *t*}, if *i* and *j* never share a roost. By choosing the parameters this way, we ensure that 0 < *γ_ij_* (*t*) < 1 for any *i* and *j*, regardless of their roosting history.

With this, the individual social influence exerted on agent *i* is no longer a constant *γ*, but an individual parameter, *γ_ij_*(*t*) ∑*_j_ γ_ij_*(*t*), that changes over time and considerably depends on the individual roosting history of an agent. Hence, the dynamics for *γ_ij_*(*t*) bridges two time scales: the time scale *t* at which agents roost together and the time scale *y* at which long-lasting social structures of the colonies, such as communities in the social network (Kerth *et al*., 2011), become visible and important. Ideally, we should observe the emergence of such communities when bridging these two time scales.

Figure 5 illustrates the impact of this model modification. The top row shows the social network of the two colonies as extracted from roost association data on the time scale *y* of a whole year (season) (Kerth *et al*., 2011). While the smaller colony, BS, does not displays any community structure, the larger colony, GB2, clearly has two communities. The bottom row of the same Figure shows the social network as obtained from our model using the adaptive *γ_ij_*(*t*). We note that these structures are observed *after* the respective *γ_ij_*(*t*) have been relaxed to some quasistationary values, i.e. after *t* = 200 days.

**Figure 5:**
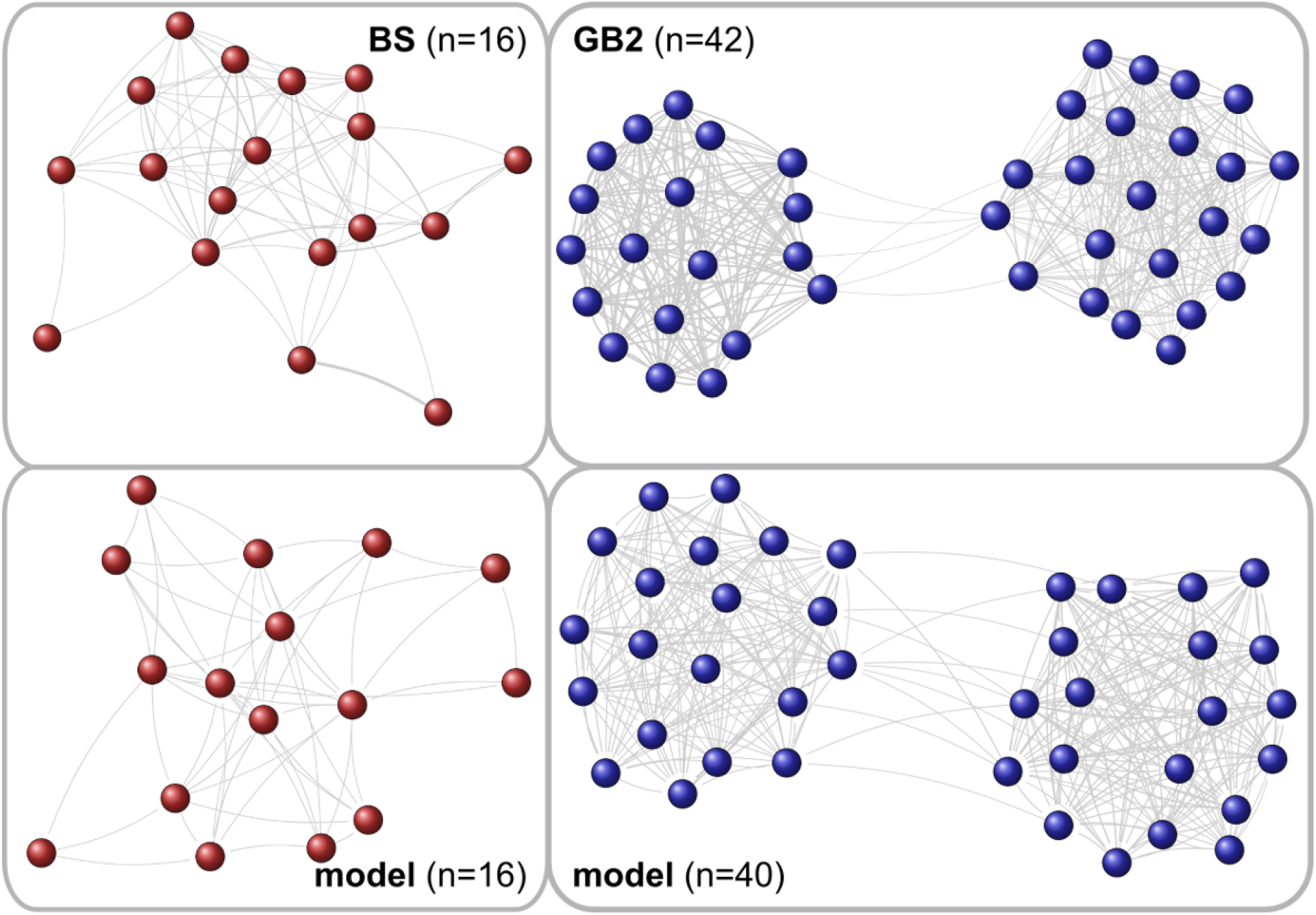
Social network of long-term roosting associations for small (left panels) and large (right panels) colonies, (top panels) Empirical data from the colonies BS (left) and GB2 (right) in the year 2007. These figures are modified from Kerth *et al*. (2011). (bottom panels) Model-generated social network with dynamic *γ_ij_* (*t*) after *t* = 200 days. For clarity, in all networks only the *strong ties*, with a weight larger than the mean value of the weight distribution, are shown.

The interesting finding here is not so much the existence of the two communities in the colony GB2. We remind that, in our initial conditions, we have already introduced two groups of size *N*/2 and have argued about slightly different initial values *γ_ij_*(0) for agents within the same group vs. agents in different groups. With this in mind, we cannot claim the *emergence* of two communities. However, we note the remarkably stable community structure: Because this colony GB2 is, with *N* = 42, well above the calculated critical colony size, *N_crit_* = 18, small initial differences in the social influences, expressed by *γ_ij_*(0), not only persisted over a long time but were amplified by the daily fission-fusion dynamics. This eventually resulted in the appearance of two separate network communities on the seasonal time scale.

The more remarkable finding is the *disappearence* of the same group structure when modeling the smaller colony BS. This colony had a size *N* = 16 *below* the critical colony size, *N_crit_* = 18. Hence, even with the same setup for *γ_ij_*(0), the daily fission-fusion dynamics was not able to sustain the induced two groups, to transform them into stable communities. Thus, on the seasonal time scale we obtain with our model the emergence of a single community that is identical with the colony. This lends strong evidence to the assumed dynamics for *γ_ij_*(*t*), and to the agent-based model of roosting behavior.

## 4 Discussion

In this paper we studied the roosting dynamics of two colonies of Bechstein’s bats, both from the *empirical* and the *modeling* perspective. Our interest was to better understand the *fission-fusion dynamics*, i.e. the formation and dissolution of different day roosting groups in these colonies. Such groups facilitate communal roosting on a daily scale, but do not form social structures stable over a long time. I.e. they are different from long-lasting community structures that can be detected in larger colonies (Kerth *et al*., 2011). In the following, we further discuss some implications of our investigations.

### Emergent structures: Group sizes, roosting durations

With our investigations, we follow a *bottom-up approach*, explaining the *emergence* of the groups from the interactions of the individuals that comprise the colonies. These interactions are described by simple rules agents follow in making roost decisions. I.e. our modeling assumptions focus on the *micro*, or agent, level, and the rules are defined on the shortest time scale *τ*. We want to reproduce the groups observed on the *macro*, or system, level on the intermediate time scale *t*.

Methodologically, we argue that our agent-based model is correct if it is able to reproduce the observed macroscopic quantities, this way explaining their emergence from micro interactions. More specifically, we can deduce that our rules are compatible with the observed system properties, that is, the size of the roosting groups. Investigations of how bats make their decisions need more data at higher temporal resolution. Hence, further research is needed to focus on this specific question. But we can clearly state that the rules that successfully describe the emergence of the system properties provide suitable *hypotheses* for the behavior of biological entities.

What kind of emergent properties can we reproduce? The first one is the exponential distribution of durations *T*, i.e. the time spent at the same roost, before switching to another roost. The quantity *T_i_* is measured for individual bats, but only the aggregation to the system level allows to determine the distribution *P*(*T*), which follows a very simple form with only one characteristic parameter *α*. Importantly, our empirical analysis shows that this distribution is independent of the colony size *N*, which also was reproduced by our model.

For the second emergent property, namely the formation of groups of different sizes inside a colony, we do have a dependence of the colony size *N*. Specifically, small colonies mostly form one roosting group, while the larger colonies mostly split into roosting groups of different sizes. This emergent behavior was also reproduced by our model.

Hence, our agent-based model is able to reproduce two very different and not directly related phenomena, which depend differently on the system size. This lends further evidence to the model.

### Modeling individual decisions

The dynamic phenomenon we are interested in basically follows a *simplified daily rhythm*. Nightly foraging, during which bats are alone or in pairs (Melber *et al*., 2013), is followed at dawn by a decision-making phase during which bats decide about the roost they will stay in for most of the following day. This leads to the formation of groups which roost together. At dusk, the bats leave their day roosts and the groups dissolve. Then, this cycle starts over again.

Our agent-based model of roosting group formation models specifically the decision-making phase occurring at dawn every day before the bats start roosting. This process is modeled at the time scale *τ*. For each agent, the roosting decision is affected by two parameters: λ describes random influences, whereas *γ* describes social influences exerted from other bats. We note that, with only the random influence, we would be able to reproduce the distribution *P*(*T*), but not the observed group structure. This leads to the conclusion that the influence coming from the roosting decisions of other bats need to be explicitly taken into account. Here we assumed that agents simply copy the roost preference of other agents at a rate *γ*. We further considered that agents finalize their roosting decisions at different times, which allows to capture the influence of agents that have already decided on those that have not decided yet.

Individual decisions balance between two concurrent requirements: the pressure to change roosts, e.g. because previous roosts are contaminated with parasites (Reckardt and Kerth, 2006), and the pressure to roost together with other colony members, for example for thermoregulatory purposes (Kerth *et al*., 2001b; Pretzlaff *et al*., 2010; Küpper *et al*., 2016). Our model reflects that information transfer between bats about roosts plays an important role. We consider that, during the decision-making phase, an individual may copy this information from another one, with a fixed rate *γ*. This way, our model presents an agent-based approach to a fully decentralised, self-organised group decision process.

The model contains two free parameters λ and *γ*. We determined these two parameters *indirectly*, by simulating the model outcome for the duration to stay in the same roost as before, which is determined by *both* parameters. We then adjusted these two parameter such that (i) the discrepancy between the observed and the modeled distribution *P*(*T*) and (ii) the variance of this distribution was minimized. We note that this *model calibration* does not involve information about the group sizes. Instead, the comparison between observed and simulated group size was used to estimate the model performance, independent of the calibration.

### Interactions in small vs. large colonies

A major empirical finding of our study was that colonies split differently into roosting groups, dependent on their size. This confirms previous empirical findings on Bechstein’s bats (Kerth and König, 1999). Small colonies mostly form one group, whereas larger colonies form several groups, the largest one comprising about half of the size of the colony. The question is whether this *transition in the system dynamics* dependent on the system size can be understood as an *emergent phenomenon*. I.e., can this transition be obtained from assuming the *same* interaction rules between agents in large and small systems, or does it imply *different* interaction rules, or rules in which an explicit size dependence is encoded? With our model, we demonstrated that this transition indeed is an emerging phenomenon that occurs at a critical system size *N_crit_*. Our simulations allowed us to determine this critical value as *N_crit_* = 18, which is also in line with observations.

Moreover, our model is able to reproduce the group sizes *both* for systems smaller and larger than *N_crit_*, using the *same interaction rules*. We found that larger colonies form groups such that the largest group comprises about one half of the colony and the second largest group about one third of the colony, which is also supported by empirical data.

### Short-term vs. long term social structures

The collective behavior of Bechstein’s bats have to be described on three different time scales. The *decisions* about roosts occurs during the swarming phase, on the time scale *τ*, shorter than one day. The *roosting in groups* occurs on the time scale *t* measured in days. I.e. every day, the groups formed before dissolve and new groups are formed. The question is how this dynamics relates to other dynamical processes observed in the colony on longer time scales. Specifically, Kerth *et al*. (2011) and Baigger *et al*. (2013) already reported that the long-term social network of larger colonies of Bechstein’s bats consists of *communities*. These are quite stable social structures that can be detected over years.

To bridge between the short-scale and the long-scale dynamics, we allowed the social influence to evolve over time, on the day time scale *t*. Specifically, we turned the homogeneous parameter, *γ*, equal for all agents, into an individual parameter *γ_i_*(*t*) = ∑_*j*_ *γ_ij_*(*t*) where *γ_ij_*(*t*) describes the mutual social influence of agents *i* and *j* as a result of their common roosting history. For the dynamics of *γ_ij_*(*t*), we adopted reinforcement learning, i.e. *γ_ij_*(*t*) increases if *i* and *j* roost together and it decreases if they don’t.

This dynamics occurs over many days, this way coupling the system dynamics on day time scales with the long-term behavior. As the result, we could demonstrate that groups existing on day time scale can translate, over time, into long-term social structures, such as communities, *if* these systems are larger than the critical size *N_crit_*. In systems smaller than *N_crit_*, on the other hand, we could show that even induced group structures cannot be transformed into long-term community structures. This agrees with empirical observations that report the absence of such community structures in small colonies (Kerth *et al*., 2011; Baigger *et al*., 2013).

We note that the daily splitting into groups and the duration of stay per roost can be modeled with a relatively simple self-organizing mechanism based on a constant *γ*. But for the formation of long-term stable communities in the larger of the two colonies, we need to introduce an individual memory effect, expressed in *γ_i_*(*t*), that allows the bats in the model to keep a record of their previous roosting history.

We conclude that our agent-based model is well posed to capture two different empirical observations in Bechstein’s bats, namely the distribution of stay lengths and the distribution of group sizes, which are not inherently connected. This lends evidence to the assumed rules of interactions because they are able to reproduce these different systemic properties. Comparing our model to other agent-based models of fission-fusion dynamics discussed above, we highlight that none so far has addressed the social and temporal aspects of the fission-fusion dynamics together. Moreover, in comparison to formally advanced, but rather abstract models (Conradt and Roper, 2000; Nair *et al*., 2019) or mere simulation approaches (Sueur and Maire, 2014), our model outcome can be directly compared to the respective empirical observations in Bechstein’s bats. This was also achieved by the agent-based model of red-capped mangabeys with respect to travel distance patters (Dolado *et al*., 2017). But their model relied on a large number of parameters proxied from field data, whereas our model has the advantage of a simple, yet convincing approach of the empirical observations.

## Acknowledgements

We thank the local forestry and conservation departments for their continuous support throughout this long-term study, and the numerous people who helped gather the field data used in this study, in particular Anja Baigger, Daniela Fleischmann and Markus Melber. This work profited strongly from the financial support of the German (DFG, KE 746/2-1/3-1/4-1/5-1/6-1) and the Swiss (SNF, 31-59556.99) national science foundations during the years covered in this study.

## Appendix

## Testing for the distribution of roosting durations

When comparing the distribution of roosting duration periods from one year to another, we found that 60% of the time for BS and 48% for GB2 the distributions were not significantly different (we compared all yearly distributions one to another; colony BS, years 2004-2008; colony GB2, years 2004-2010; two-sample Kolmogorov-Smirnov test with Šidák correction for multiple sampling, *n* = 10 for BS and *n* = 21 for GB2, *p* > 0.05.

Finally, the aggregated distributions of roosting duration periods for each colony did not differ significantly from one another, neither in their exponential fit (Fig. 1) nor in their observed distribution (colony BS, years 2004-2008: *n* = 3633 roosting periods in total; colony GB2, years 2004-2010: *n* = 10385 two-sample Kolmogorov-Smirnov test, *p* = 0.63).

Based on these observations, we concluded that the durations to stay in a given roost varied neither within the colonies, with different years of observation (hence, different colony sizes), nor between the two observed colonies (despite them having distinct roosting areas, see for example Kerth and van Schaik, 2012).

## Testing for the influence of distances

For each bat in each year we randomised the sequence of visited roosts and compared the distribution of flying distances aggregated over a large number of randomisations to the observed distribution (a Monte-Carlo simulation of the roosting sequence).

We found that 91% of the time, the sequence of distances between consecutively occupied roosts for a bat was not significantly different from what it would be if the bat had visited the same roosts in a random order (colony GB2, years 2005-2010; average number of roosts successively occupied per individual and per year, *n* = 38.8 ± 12.3; non-parametric two-sample Kolmogorov-Smirnov test with Šidák correction for multiple sampling within years, *n* = 231 individuals over all 5 years, *p* > 0.05 for 211 out of 231 series of observations).

Moreover, we identified no effect of the previously occupied roost on the next occupied roost in terms of flying distance between roosting sites. In light of these observations, and in order to adopt a parsimonious approach, we modelled in a first step the roosting behaviour of an individual bat by a zero-order Markov process (or random walk).

## References

Anderson, P. (1972). More is different. Science 177(4047), 393–396.

Aureli, F.; Shaffner, C.; Boesch, C.; Bearder, S.; Call, J.; Chapman, C.; Connor, R.; Di Fiore, A.; Dunbar, R.; Henzi, S.; et al. (2008). Fission-fusion dynamics: new research frameworks. Current Anthropology 49(4), 627–654.

Baigger, A.; Perony, N.; Leinert, V.; Melber, M.; Grunberger, S.; Fleischmann, D.; Kerth, G. (2013). Bechstein’s bats maintain individual social links despite a complete reorganisation of their colony structure. Naturwissenschaften 100(9), 895–898.

Body, G.; Weladji, R. B.; Holand, O.; Nieminen, M. (2015). Measuring variation in the frequency of group fission and fusion from continuous monitoring of group sizes. Journal of Mammalogy 96(4), 791–799.

Bonabeau, E.; Dagorn, L. (1995). Possible universality in the size distribution of fish schools. Physical Review E 51(6), 5220–5223.

Bonabeau, E.; Dagorn, L.; Fréon, P. (1999). Scaling in animal group-size distributions. Proceedings of the National Academy of Sciences 96(8), 4472.

Buhl, J.; Sumpter, D.; Couzin, I.; Hale, J.; Despland, E.; Miller, E.; Simpson, S. (2006). From disorder to order in marching locusts. Science 312(5778), 1402.

Cavagna, A.; Giardina, I. (2010). Large-scale behaviour in animal groups. Behavioural Processes 84(3), 653–656.

Conradt, L.; Roper, T. (2005). Consensus decision making in animals. Trends in Ecology & Evolution 20(8), 449–56.

Conradt, L.; Roper, T. (2007). Democracy in animals: the evolution of shared group decisions. Proceedings of the Royal Society B: Biological Sciences 274(1623), 2317.

Conradt, L.; Roper, T. J. (2000). Activity synchrony and social cohesion: a fission-fusion model. Proceedings of the Royal Society of London. Series B: Biological Sciences 267(1458), 2213–2218.

Conradt, L.; Roper, T. J. (2003). Group decision-making in animals. Nature 421(6919), 155–8.

Couzin, I. (2009). Collective cognition in animal groups. Trends in Cognitive Sciences 13(1), 36–43.

Couzin, I.; Krause, J. (2003). Self-organization and collective behavior in vertebrates. Advances in the Study of Behavior 32, 1–75.

Couzin, I.; Laidre, M. (2009). Fission-fusion populations. Current Biology 19(15), R633.

Dolado, R.; Gimeno, E.; Beltran, F. S. (2017). Modeling the emergence of seasonal fission-fusion dynamics in red-capped mangabeys (Cercocebus torquatus). Behavioral Ecology and Sociobiology 71(7), 100.

Entwistle, A.; Racey, P.; Speakman, J. (1996). Habitat exploitation by a gleaning bat, Plecotus auritus. Philosophical Transactions of the Royal Society of London. Series B: Biological Sciences 351(1342), 921–931.

Fleischer, T.; Gampe, J.; Scheuerlein, A.; Kerth, G. (2017). Rare catastrophic events drive population dynamics in a bat species with negligible senescence. Scientific Reports 7, 7370.

Fleischmann, D.; Baumgartner, I. O.; Erasmy, M.; Gries, N.; Melber, M.; Leinert, V.; Parchem, M.; Reuter, M.; Schaer, P.; Stauffer, S.; Wagner, I.; Kerth, G. (2013). Female Bechstein’s Bats Adjust Their Group Decisions about Communal Roosts to the Level of Conflict of Interests. Current Biology 23(17), 1658–1662.

Fleischmann, D.; Kerth, G. (2014). Roosting behavior and group decision making in 2 syntopic bat species with fission-fusion societies. Behavioral Ecology 25(5), 1240–1247.

Flierl, G.; Grünbaum, D.; Levins, S.; Olson, D. (1999). From individuals to aggregations: the interplay between behavior and physics. Journal of Theoretical Biology 196(4), 397–454.

Gautrais, J.; Theraulaz, G.; Deneubourg, J.; Anderson, C. (2002). Emergent polyethism as a consequence of increased colony size in insect societies. Journal of Theoretical Biology 215(3), 363–373.

Jeanson, R.; Fewell, J.; Gorelick, R.; Bertram, S. (2007). Emergence of increased division of labor as a function of group size. Behavioral Ecology and Sociobiology 62(2), 289–298.

Kerth, G. (2006). Relatedness, life history and social behaviour in the long-lived Bechstein’s bat (Myotis bechsteinii). In: Z. Akbar; G. McCracken; T. Kunz (eds.), Functional and evolutionary ecology of bats, Oxford University Press, pp. 199–212.

Kerth, G. (2008). Causes and Consequences of Sociality in Bats. BioScience 58(8), 737.

Kerth, G.; Ebert, C.; Schmidtke, C. (2006). Group decision making in fission-fusion societies: evidence from two-field experiments in Bechstein’s bats. Proceedings of the Royal Society B: Biological Sciences 273(1602), 2785.

Kerth, G.; König, B. (1999). Fission, fusion and nonrandom associations in female Bechstein’s bats (Myotis bechsteinii). Behaviour 136(9), 1187–1202.

Kerth, G.; Perony, N.; Schweitzer, F. (2011). Bats are able to maintain long-term social relationships despite the high fission-fusion dynamics of their groups. Proceedings of the Royal Society B: Biological Sciences 278(1719), 2761–2767.

Kerth, G.; Reckardt, K. (2003). Information transfer about roosts in female Bechstein’s bats: an experimental field study. Proceedings of the Royal Society B: Biological Sciences 270(1514), 511.

Kerth, G.; van Schaik, J. (2012). Causes and consequences of living in closed societies: lessons from a long-term socio-genetic study on Bechstein’s bats. Molecular Ecology 21(3), 633–646.

Kerth, G.; Wagner, M.; König, B. (2001a). Roosting together, foraging apart: information transfer about food is unlikely to explain sociality in female Bechstein’s bats (Myotis bechsteinii). Behavioral Ecology and Sociobiology 50(3), 283–291.

Kerth, G.; Weissmann, K.; König, B. (2001b). Day roost selection in female Bechstein’s bats (Myotis bechsteinii): a field experiment to determine the influence of roost temperature. Oecologia 126(1), 1–9.

Krause, J.; Ruxton, G. (2002). Living in groups. Oxford University Press.

Küpper, N. D.; Melber, M.; Kerth, G. (2016). Nightly clustering in communal roosts and the regular presence of adult females at night provide thermal benefits for juvenile Bechstein’s bats. Mammalian Biology - Zeitschrift für Säugetierkunde 81(2), 201–204.

Melber, M.; Fleischmann, D.; Kerth, G. (2013). Female Bechstein’s Bats Share Foraging Sites with Maternal Kin but do not Forage Together with them–Results from a Long-Term Study. Ethology 119(9), 793–801.

Nair, G. G.; Senthilnathan, A.; Iyer, S. K.; Guttal, V. (2019). Fission-fusion dynamics and group-size-dependent composition in heterogeneous populations. Physical Review E 99(3), 032412.

Nad’o, L.; Kaňuch, P. (2015). Swarming behaviour associated with group cohesion in tree-dwelling bats. Behavioural processes 120, 80—86.

O’Donnell, C. F.; Sedgeley, J. A. (1999). Use of roosts by the long-tailed bat, Chalinolobus tuberculatus, in temperate rainforest in New Zealand. Journal of Mammalogy 80(3), 913—923.

Okubo, A. (1986). Dynamical aspects of animal grouping: swarms, schools, flocks, and herds. Advances in Biophysics 22, 1–94.

Pretzlaff, I.; Kerth, G.; Dausmann, K. (2010). Communally breeding bats use physiological and behavioural adjustments to optimise daily energy expenditure. Naturwissenschaften 97(4), 353–363.

Ramos-Fernández, G.; Boyer, D.; Gómez, V. (2006). A complex social structure with fission-fusion properties can emerge from a simple foraging model. Behavioral Ecology and Sociobiology 60(4), 536–549.

Ramos-Fernández, G.; Morales, J. M. (2014). Unraveling fission-fusion dynamics: How subgroup properties and dyadic interactions influence individual decisions. Behavioral Ecology and Sociobiology 68(8), 1225–1235.

Reckardt, K.; Kerth, G. (2006). The reproductive success of the parasitic bat fly Basilia nana (Diptera: Nycteribiidae) is affected by the low roost fidelity of its host, the Bechstein’s bat (Myotis bechsteinii). Parasitology Research 98(3), 237–243.

Sueur, C.; King, A.; Conradt, L.; Kerth, G.; Lusseau, D.; Mettke-Hofmann, C.; Schaffner, C.; Williams, L.; Zinner, D.; Aureli, F. (2011). Collective decision-making and fission-fusion dynamics: a conceptual framework. Oikos 120, 1608–1617.

Sueur, C.; Maire, A. (2014). Modelling animal group fission using social network dynamics. PLoS ONE 9(5), 0097813.

Sumpter, D. (2010). Collective animal behavior. Princeton University Press.

Van Horn, R.; Buchan, J.; Altmann, J.; Alberts, S. (2007). Divided destinies: group choice by female savannah baboons during social group fission. Behavioral Ecology and Sociobiology 61(12), 1823–1837.

Willis, C.; Brigham, R. (2004). Roost switching, roost sharing and social cohesion: forest-dwelling big brown bats, Eptesicus fuscus, conform to the fission-fusion model. Animal Behaviour 68(3), 495–505

